# The Immune-Evasive Proline 283 Substitution in Influenza Nucleoprotein Increases Aggregation Propensity Without Altering the Native Structure

**DOI:** 10.1101/2023.09.08.556894

**Authors:** Jimin Yoon, Yu Meng Zhang, Cheenou Her, Robert A. Grant, Anna M. Ponomarenko, Bryce E. Ackermann, Galia T. Debelouchina, Matthew D. Shoulders

## Abstract

Nucleoprotein (NP) is a key structural protein of influenza ribonucleoprotein complexes and is central to viral RNA packing and trafficking. In human cells, the interferon induced Myxovirus resistance protein 1 (MxA) binds to NP and restricts influenza replication. This selection pressure has caused NP to evolve a few critical MxA-resistant mutations, particularly the highly conserved Pro283 substitution. Previous work showed that this essential Pro283 substitution impairs influenza growth, and the fitness defect becomes particularly prominent at febrile temperature (39 °C) when host chaperones are depleted. Here, we biophysically characterize Pro283 NP and Ser283 NP to test if the fitness defect is owing to Pro283 substitution introducing folding defects. We show that the Pro283 substitution changes the folding pathway of NP without altering the native structure, making NP more aggregation prone during folding. These findings suggest that influenza has evolved to hijack host chaperones to promote the folding of otherwise biophysically incompetent viral proteins that enable innate immune system escape.

**Teaser:** Pro283 substitution in flu nucleoprotein introduces folding defects, and makes influenza uniquely dependent on host chaperones.

## Introduction

Influenza is a pathogenic RNA virus that causes acute respiratory disease in humans. The influenza genome consists of eight segments of negative-sense single-stranded RNA, with each segment assembled into a double-helical viral ribonucleoprotein (*1–3*). Each ribonucleoprotein complex is composed of one strand of viral RNA encapsidated by nucleoprotein (NP) molecules and three viral polymerase subunits.

A critical structural component of the ribonucleoprotein complex, NP is a 56 kDa globular protein with key roles in viral RNA packing and trafficking (*4*), transcription and replication (*5*) and progeny virion assembly (*6*). Like many other viral proteins that are recognized by cellular immune factors, NP is mainly targeted by the interferon-induced Myxovirus resistance protein 1 in human cells (MxA) (*7, 8*). MxA has been proposed to bind to NP and self-assemble into a higher-order oligomeric complex, restricting influenza viral ribonucleoprotein access to the host nucleus and potentially also preventing proper vRNP assembly (*9–11*). Evading the antiviral activity of MxA is crucial for the continued pathogenicity of the influenza virus. The imperative to escape this selection pressure has caused NP to acquire a number of MxA-resistant mutations (*7*).

The Pro283 substitution is one of three critical MxA-resistant substitutions that were first identified in the pandemic strain A/Brevig Mission/1918 (H1N1) (*7, 12*). Pro283 is now almost universally conserved in the circulating human isolates representing the H1N1, H1N2, H2N2, and H3N2 influenza subtypes. In contrast, the substitution is rarely found in avian influenza strains including H5N1 viruses, where the avian MxA ortholog lacks potency against influenza (*7, 12–14*). Nonetheless, while allowing for MxA escape, Pro283 substitution impairs influenza growth when introduced to H5N1 or H7N7 viruses (*7, 15*). Furthermore, a more recent study found that the fitness of Pro283 NP-encoding influenza strongly depends on both temperature and the availability of host cell chaperones (*13*). In this study, Pro283 NP-encoding influenza was competed with influenza encoding other amino acids at site 283 of NP, such as Ser283. The relative fitness of Pro283 NP-encoding influenza was observed to be substantially compromised at a febrile temperature. The fitness decreases even further when HSF1-regulated host chaperones are depleted.

Together, these studies demonstrate that Pro283, despite being an important immune-escape substitution, has substantial associated fitness costs. The observations that the viability of Pro283-encoding influenza depends on both temperature and the availability of HSF1-regulated chaperones suggest that the fitness defect can potentially be attributed to protein-folding challenges caused by the Pro283 substitution, yet no study has identified a molecular-level explanation for this fitness defect. Elucidating the underlying molecular basis of fitness loss caused by the Pro283 substitution would yield key insights into the molecular-level constraints on viral protein evolution, and how host cell chaperones may be hijacked to address these constraints and expand viral mutational landscapes (*16*).

In this work, we investigated the structural and folding consequences of the Pro283 substitution using a comprehensive suite of biophysical techniques. We observed that Pro283 NP exhibits lower thermal stability than Ser283 NP both in test tubes and in cellular environments. Perhaps surprisingly, our crystal structures and nuclear magnetic resonance (NMR) spectra revealed that the proline substitution, despite being located in the middle of an α-helix, does not substantially change the structure of NP’s folded state. However, the two NP variants did display distinct native-state dynamics in solution, with Ser283 NP exhibiting more motions in the μs–ms timescale. Most importantly, unfolding/refolding experiments showed that Pro283 NP can undergo a three-state unfolding transition, whereas Ser283 NP exhibits a two-state unfolding transition. The intermediate conformer only observed in Pro283 NP was aggregation-prone, preventing spontaneous refolding of Pro283 NP. Overall, our results indicate that Pro283 renders NP more aggregation-prone during folding, without inducing major perturbations to the final structure of the protein. Host proteostasis factors, particularly chaperones, likely mitigate this biophysical defect by preventing the aggregation of Pro283 NP and/or altering the folding pathway traversed by the nascent protein.

## Results

### Pro283 NP denatures and aggregates at a lower temperature than Ser283 NP

To probe the biophysical consequences of a proline residue at site 283 in an otherwise identical protein sequence background, we began with the A/Aichi/1968 (H3N2) influenza A nucleoprotein that has wild-type Pro283 and constructed a Pro283Ser variant-encoding plasmid. We chose Ser283 NP for comparison because Ser283 is the wild-type amino acid for the well-characterized A/WSN/33 (H1N1) strain of influenza, and it was previously used in head-to-head competition experiments that revealed the temperature and host-chaperone dependence of Pro283 NP-encoding influenza. We recombinantly expressed and purified His_6_-tagged Pro283 and Ser283 variants from *Escherichia coli* without noticeable yield differences. Since NP binds to RNA in cells, it expresses as monomers and oligomers that exist in a dynamic equilibrium (*17*). To isolate monomeric NP, which is important for biophysical techniques that require a homogenous sample, we used a NP construct previously engineered to lack RNA binding ability that includes the R416A substitution and deletion of residues 2–7 (*13, 17–19*). We used size exclusion chromatography to show that both Pro283 and Ser283 were isolated as pure monomers (**fig. S1A** and **S1B**). We further verified the monomeric assembly state via dynamic light scattering and sedimentation velocity experiments (**fig. S1C** and **S1D**). Both experiments showed that >97% of the protein species in the sample were monomeric NP (actual molecular weight = 56 kDa).

Since previous deep mutational scanning data show that the viability of influenza virus carrying Pro283 NP is compromised at 39 °C (*13*), we first tested whether purified Pro283 also exhibited lower thermal stability compared to Ser283 NP using differential scanning fluorometry (*20*). The SYPRO Orange fluorescence spectra showed that Pro283 NP exhibited higher initial fluorescence and a smaller increase in fluorescence upon heating, potentially suggesting that Pro283 NP has more exposed hydrophobic residues in its native state (**Fig. 1A**). Circular dichroism spectra of the two variants were nonetheless indistinguishable, suggesting that the secondary structure content was generally similar between the two variants (**Fig. 1B**). The fluorescence for both variants decreased after reaching a peak, likely indicating that NP is aggregating after the melting temperature that eliminates available hydrophobic residues that can interact with SYPRO Orange (*21*). Since protein aggregation is often concentration-dependent, we performed thermal melts at two different protein concentrations. We fit the curves to a two-state unfolding and observed that *T*_agg_ of Pro283 NP is ∼2°C lower than *T*_agg_ of Ser283 NP at all protein concentrations tested (**Fig. 1C**). This result agrees with a previous circular dichroism-based thermal aggregation experiment that showed Pro283 irreversibly precipitating at 40.3 °C, while Ser283 precipitated at 43.0 °C, leading to complete signal loss (*22*). Further, we observed that the difference in the *T*_agg_ values of the two variants increased with decreasing protein concentration (*T*_agg_ of Ser283 NP − *T*_agg_ Pro283 NP = 1.6 °C at 1 mg/mL, 2.1 °C at 0.2 mg/mL, and 2.7 °C at 0.05 mg/mL).

**Figure 1.**
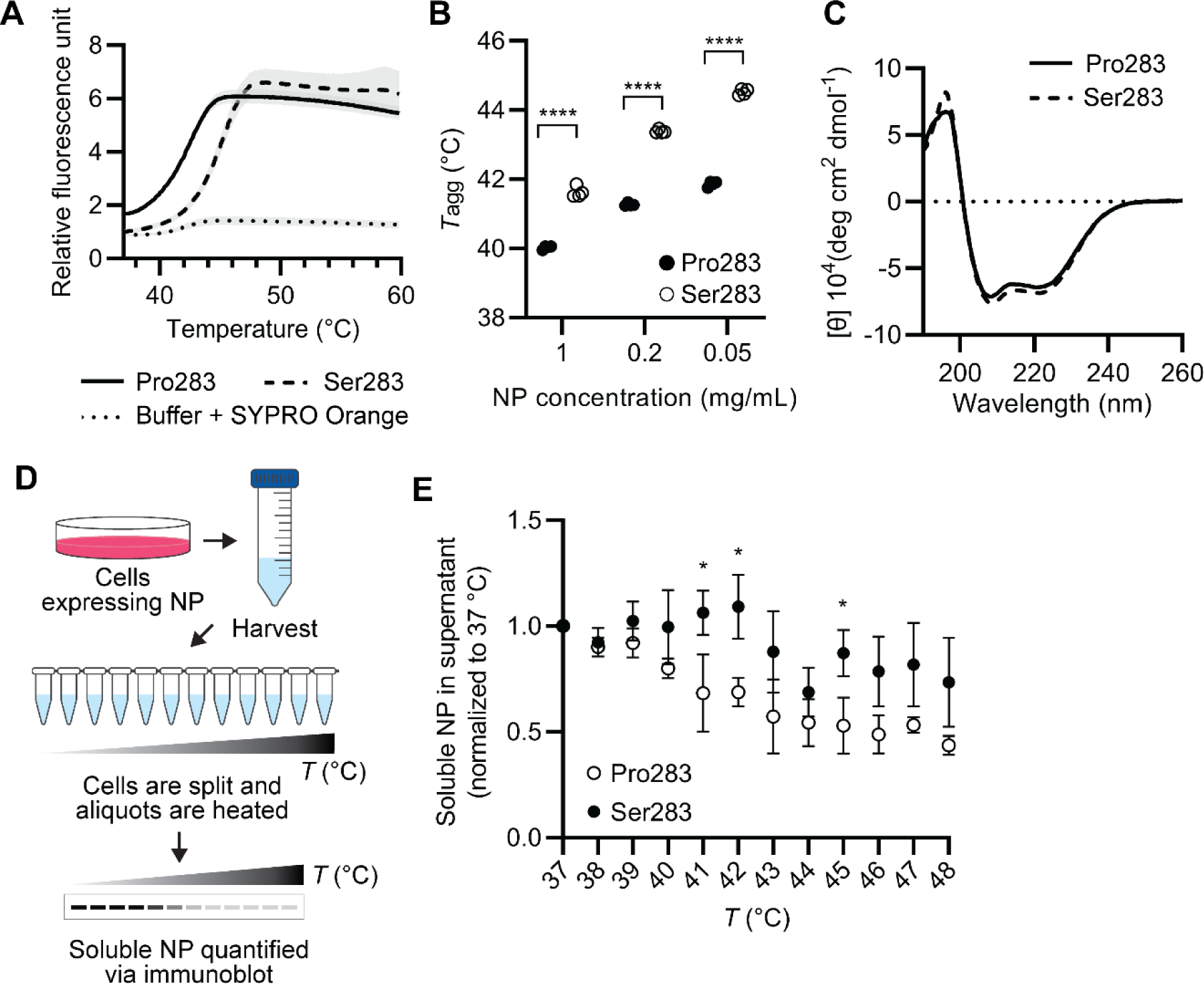
Pro283 NP has lower thermal stability than Ser283 NP both in test tube and in cells. (**A**) Representative SYPRO Orange thermal unfolding curve of Pro283 NP, Ser283 NP, and SYPRO Orange dye with buffer only. Mean of four technical replicates are shown with the shaded area representing the standard deviation. (**B**) Irreversible aggregation temperature (*T*_agg_) was calculated by fitting the plots from (**A**) to a Boltzmann sigmoidal curve. The experiment was performed at three different NP concentrations. Individual measurements for five technical replicates are shown (**** indicates *p*-value <0.0001). (**C**) CD spectra of Pro283 and Ser283 NP (**D**) Schematic for a cellular thermal shift assay experiment. Intact cells ectopically expressing NP are harvested, split into aliquots, and heated rapidly to the indicated temperatures in a thermocycler. After centrifugation to pellet protein aggregates, soluble NP remaining in the supernatant was quantified via immunoblot. (**E**) Quantification of cellular thermal shift assay immunoblots. The mean of biological triplicates is shown with the error bars representing standard deviations (* indicates *p*-value < 0.05). All band intensities were normalized to 37 °C. Original blots are shown in **fig. S2**.

While these results indicate that Pro283 NP has a lower thermal stability in a test tube, the result could be different in cells. To test if Pro283 exhibits lower thermal stability in living cells, we performed a cellular thermal shift assay (*23*). Briefly, cells ectopically expressing the Pro283 or Ser283 NP with C-terminal HA tags replacing the His_6_ tags were rapidly heated to promote irreversible protein aggregation, and the amount of NP remaining in the soluble fraction of cell lysates was analyzed via immunoblot (**Fig. 1D**). The NP construct used for cellular thermal shift assay also did not have the R416A substitution and 2–7 deletion that enforced monomerization, better representing the behavior of NP in cells. Overall, the aggregation curves for both variants did not fit well to a two-state model, making *T*_agg_ determination difficult. Nevertheless, it was clear that Pro283 NP began aggregating at lower temperatures than Ser283, as can be seen by the lower fraction of soluble NP remaining at 41 °C and 42 °C for Pro283. (**Fig. 1E** and **fig. S2**). Aggregation of Ser283 did not initiate until ∼43 °C. Together, these data show that Pro283 NP has lower thermal stability than Ser283 NP both in test tubes and in cellular environments.

### Pro283 substitution does not substantively alter the native structure of NP

Structures of other influenza NP homologs, mostly from avian influenza strains, have a serine or leucine at site 283, and reveal that site 283 lies in the middle of an α-helix (*17, 24*). Since proline is known to disrupt α-helical conformations (*25*), we initially hypothesized that this proline substitution would disrupt the local secondary structure, leading to the observed lower thermal robustness and chaperone dependence.

To test this hypothesis, we obtained high-resolution X-ray crystal structures of Pro283 and Ser283 NP to 2.90 Å and 3.09 Å resolution, respectively. Both structures were solved via molecular replacement using a monomeric nucleoprotein of A/WSN/1933 (H1N1) influenza (PDBID: 3ZDP) (*19*) as the search model, and both structures belonged to the space group C222_1_. The final models for both variants contained three molecules per asymmetric unit. The molecules within an asymmetric unit had the same conformations with low root mean square deviations (RMSDs) (**fig. S3**), Chain A was used for subsequent analyses for both models. There was no density observed for residues 1–20, 401–412 and 454–457 so these residues were not included in the final models. The crystallization conditions are reported in **Table 1** and data and refinement statistics are reported in **Table 2**.

**Table 1.**
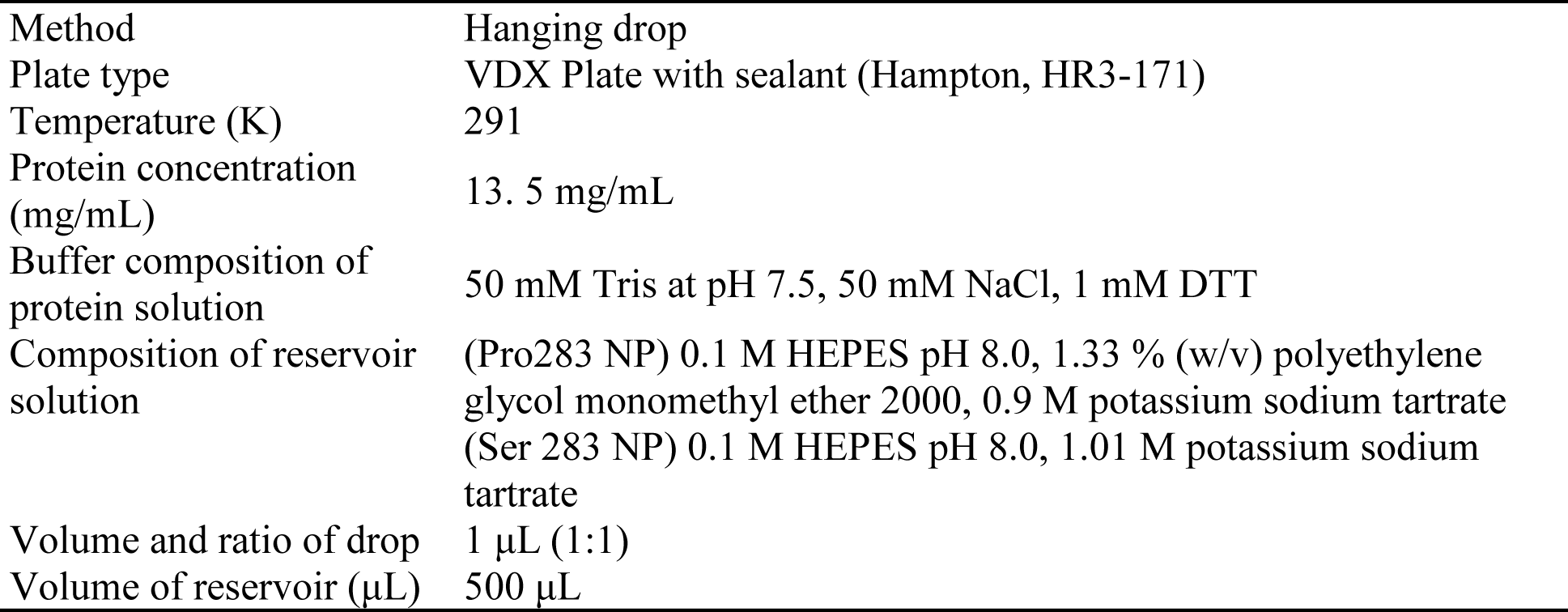
NP crystallization conditions.

**Table 2.** X-ray crystallography data statistics.

|  | Pro283 | Ser283 |
| --- | --- | --- |
| Resolution limit (Å) | 48.8–2.90 (2.98–2.90) | 48.9–3.09 (3.26–3.09) |
| Space group | C222<sub>1</sub> | C222<sub>1</sub> |
| Unit cell dimensions (Å) | 163.84, 282.92, 116.14 | 163.42, 285.22, 116.90 |
| $\alpha$, $\beta$, $\gamma$ (°) | 90.000, 90.000, 90.000 | 90.000, 90.000, 90.000 |
| Total No. of reflections | 321749 (20810) | 427717 (25333) |
| No. of unique reflections | 59023 (4122) | 50131 (3503) |
| R<sub>meas</sub> (%) | 9.0 (138.8) | 15.1 (142.3) |
| R<sub>merge</sub> (%) | 8.1 (124.4) | 14.2 (132.3) |
| $\langle I/\sigma \rangle$ | 15.71 (1.84) | 12.76 (2.14) |
| CC<sub>1/2</sub> | 99.9 (64.6) | 99.8 (69.0) |
| Completeness (%) | 98.1 (94.2) | 99.6 (96.6) |
| Redundancy | 5.5 (5.0) | 8.5 (7.2) |
| Overall B factor from Wilson plot (Å<sup>2</sup>) | 83.92 | 93.88 |
| Reflections used in refinement | 59022 (5598) | 50081 (4786) |
| Reflections used for R-free | 1995 (190) | 2005 (191) |
| R<sub>work</sub> | 0.2198 (0.3657) | 0.2202 (0.3992) |
| R<sub>free</sub> | 0.2445 (0.3979) | 0.2355 (0.4290) |
| No. of non-hydrogen atoms |  |  |
| Protein | 10734 | 10737 |
| Ligands | 0 | 1 |
| Solvent | 0 | 0 |
| RMSD from ideal values |  |  |
| Bond length (Å) | 0.002 | 0.003 |
| Bond angle (°) | 0.56 | 0.59 |
| Ramachandran plot |  |  |
| Most favored (%) | 98.43 | 97.84 |
| Allowed (%) | 1.57 | 2.08 |
| Outlier (%) | 0 | 0.07 |
| Rotamer outliers (%) | 0.26 | 0.53 |
| Average B factors |  |  |
| Protein atoms | 102.24 | 97.44 |
| Ligand atoms |  | 73.24 |

\*Highest resolution bin statistics indicated in parentheses.

Our crystal structures revealed that the Pro283 substitution disrupts neither the local secondary structure nor the gross tertiary structure. The Pro283 and Ser283 NP structures were highly similar, with an overall RMSD of only 0.347 Å (**Fig. 2A**). For both structures, site 283 was located at the center of a two-turn α-helix spanning residues Ala278–Ser287. Surprisingly, introduction of Pro283 into the middle of α-helix barely perturbed the helix, and the main chain traces of the Pro283 α-helix and Ser283 α-helix had an RMSD of only 0.399 Å (**Fig. 2B**). This conservation of the α-helix despite the presence of proline was recently independently confirmed by a crystal structure of A/Northern Territory/1968 (H3N2) influenza NP (PDBID: 7NT8), which was not available at the time we were pursuing our structures (*26*). The Pro283 NP structure was also very similar to the previously reported H1N1 NP (PDBID: 3ZDP, RMSD: 0.490 Å) structure, and to the H5N1 NP (PDBID: 2Q06, RMSD: 0.793 Å) structure except for the C-terminal disordered regions, which had serine and leucine at position 283, respectively.

**Figure 2.**
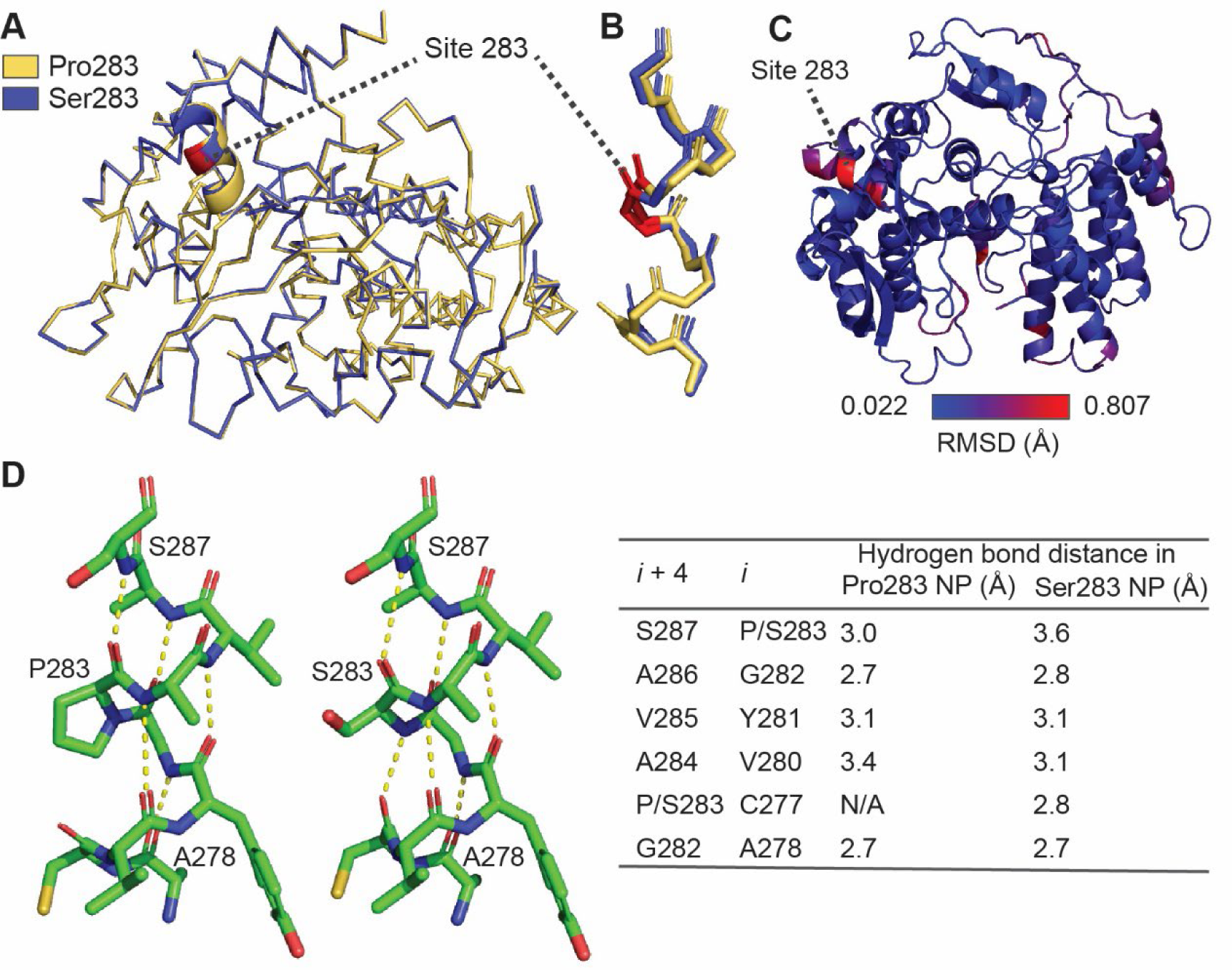
X-ray crystal structures of Pro283 and Ser283 NP are highly similar. (**A**) Overlay of the C_α_ traces from the Pro283 and Ser283 NP crystal structures. Site 283 is marked as red. Yellow: Pro283 NP; blue: Ser283 NP. (**B**) Overlay of the main chain atoms for the α-helix containing site 283 (spanning Ala278 to Ser287). Site 283 is marked as red. Yellow: Pro283 NP; blue: Ser283 NP. (**C**) RMSD per residue mapped onto the crystal structure of Ser283 NP. Site 283 is marked with an arrow. (**D**) Stick representation of all atoms in the α-helix containing site 283 (Ala278 to Ser287), with the predicted *i* + 4 → *i* hydrogen bonds shown as yellow dashed lines. Select residues are indicated for orientation clarity. The table shows all possible *i* + 4 → *i* hydrogen bonds within the α-helix containing site 283 (Ala278–Ser287) and their distances in each of the crystal structure.

Nevertheless, the Ser283 and Pro283 NP structures were not entirely identical. First, when we calculated RMSD per residue, we observed that residues with RMSD >0.4 Å were most heavily clustered in the helix containing site 283 (**Fig. 2C**). Site 283 itself had the highest RMSD of the entire structure (0.807 Å). The absence of an amide hydrogen on Pro283 results in the loss of the 283 → Cys279 hydrogen bond (**Fig. 2D**). The remaining hydrogen bond distances for *i* + 4 amide hydrogen → *i* carbonyl oxygen were comparable between the two structures, except that the Ser287 → Pro/Ser283 distance was extended by 0.6 Å in the Ser283 structure, likely weakening that hydrogen bond. In the Pro283 NP structure, the *ψ* angle for Cys279 and the *φ* angle for Val280 also changed moderately (<35 °) to prevent a steric clash between the carbonyl oxygen of Cys279 and the δ-carbon of Pro283, but the rest of the backbone dihedral angles were largely unchanged (**fig. S4**). In sum, although we observed subtle structural differences between Pro283 and Ser283 NP in the single crystals, the differences appeared to be driven by accommodating the requirements of the pyrrolidine ring, and none were substantial enough to entirely disrupt the local α-helix or globally alter the tertiary structure in the solid-state.

### Pro283 substitution alters the dynamics of native-state NP in solution

While X-ray crystallography provides rich, atomic-resolution data on protein structures, it only captures a single conformation most favored in the context of crystal packing. The reported temperature stability differences between Pro283 and Ser283, on the other hand, may arise from variations in their dynamic properties and/or conformational ensembles in solution. To gain insights into these properties, we characterized both variants by nuclear magnetic resonance (NMR) spectroscopy. First, we assessed the stability of the proteins in buffer and concentration conditions suitable for NMR experiments at 25 °C and 37 °C (**fig. S5A** and **S5B**). At 25 °C, the ^1^H intensity of the amide region was unchanged over the course of five days at NP concentrations of 190–250 μM, suggesting that the proteins remained stable and the samples were suitable for collection of multidimensional experiments. At 37 °C, on the other hand, visible precipitation was observed and the ^1^H signals significantly diminished in intensity (**fig. S5C** and **S5D**). We therefore performed all subsequent NMR experiments at 25 °C.

We next expressed ^2^H,^13^C,^15^N-lableed NP and pursued 2D ^1^H-^15^N transverse relaxation-optimized spectroscopy heteronuclear single quantum coherence (TROSY HSQC), 3D HNCACB, and HN(CO)CACB experiments. Unfortunately, even with deuteration, there was still substantial signal overlap and ambiguity, along with low sensitivity in the 3D experiments, which precluded sequential assignments. Nonetheless, the ^1^H-^15^N TROSY HSQC provided a number of important insights. First, both variants displayed well dispersed spectra with chemical shifts consistent with a folded structure rich in α-helical content (**Fig. 3A**). Second, analysis of the spectra yielded ∼260 identifiable non-sidechain peaks for the Ser283 sample (∼52% of the sequence), and ∼300 peaks for the Pro283 variant (∼60% of the sequence). Based on their unique and well dispersed chemical shifts, we were also able to identify 26/39 glycine and 15/65 serine/threonine residues in the Pro283 NP and 24/39 glycine and 13/66 serine/threonine residues in the Ser283 NP. The substantial number of missing residues suggests that both variants contain regions with intermediate exchange dynamics that are invisible in the NMR experiments. Still, such motions (typically in the μs-ms regime) were more prevalent in the Ser283 variant at 25 °C. Third, qualitative comparison of the chemical shifts for the resolved cross-peaks in the two spectra showed a high degree of overlap, suggesting that the two variants have grossly similar structures (**Fig. 3B-D**). There are, however, some peak shifts that indicated local perturbations in the chemical environment. These observations agree with the conclusions from the crystal structures, which suggest that the structural variation between the two proteins is relatively subtle. In solution, however, we detected differences in the μs-ms dynamics between the two variants.

**Figure 3.**
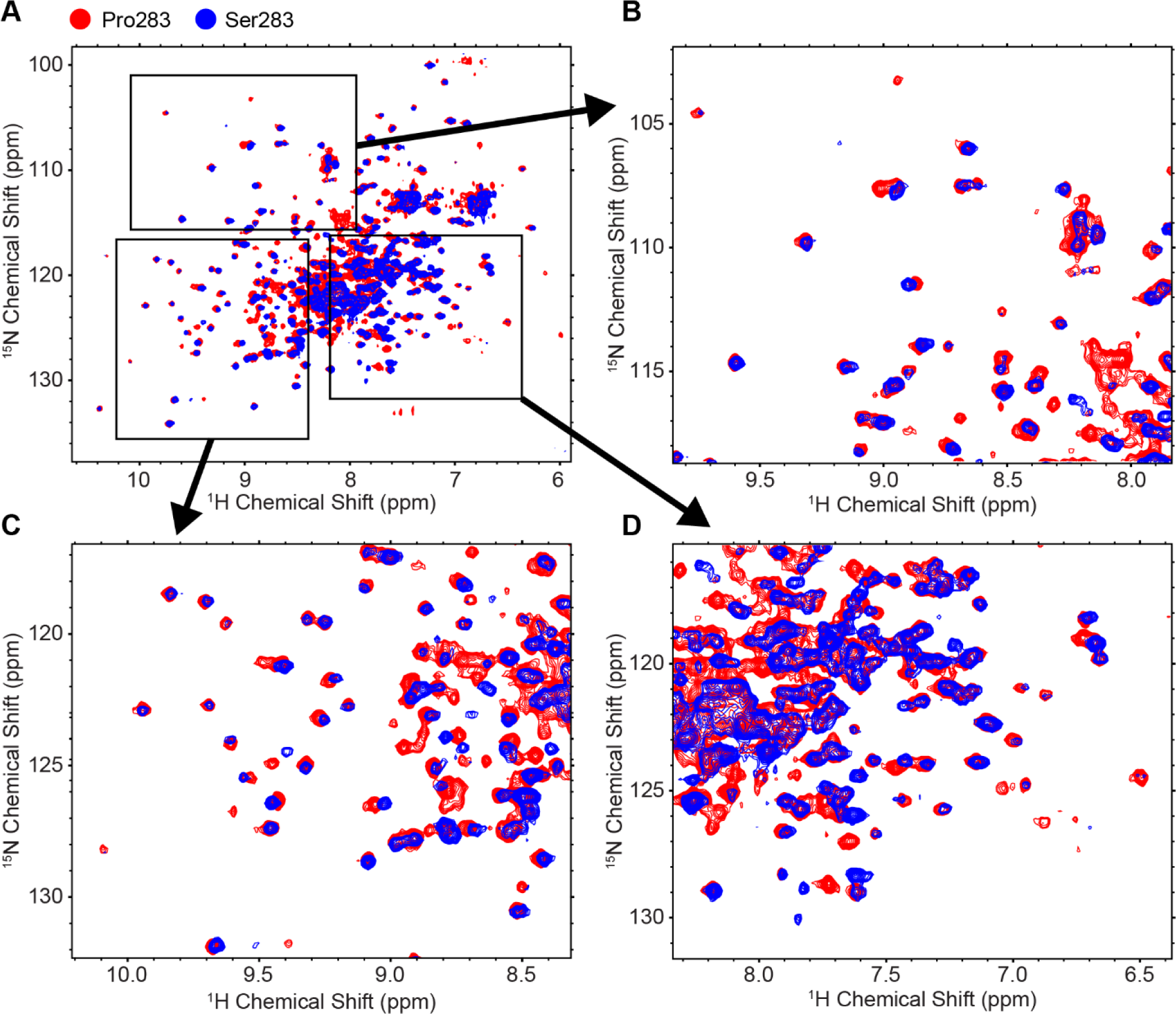
Pro283 and Ser283 NP have similar structures but distinct dynamics. Overlay of 2D ^1^H-^15^N HSQC TROSY experiments of Pro283 (red) and Ser283 (blue) NP at 25 °C.

### Pro283 substitution changes the folding pathway of NP by introducing an intermediate conformer

While Pro283 NP is MxA-evasive, it still supports viral ribonucleoprotein assembly and influenza replication, consistent with the notion that the Pro283 substitution should not drastically alter the functional conformation of native NP. Nevertheless, a substitution that does not alter the native-state conformation of a protein can still introduce biophysical defects by altering the folding pathway – for example, by slowing down the rate of folding and/or introducing a defective intermediate conformation. Therefore, we next hypothesized that the Pro283 substitution has biophysically deleterious consequences that are mediated by altering the folding pathways of NP rather than by dramatically changing NP’s native state.

To characterize the folding of Pro283 versus Ser283 NP, we performed a series of equilibrium unfolding and refolding experiments at room temperature. Upon denaturation with 6 M guanidine hydrochloride (GdnHCl), the intrinsic fluorescence of NP changed substantially for both Pro283 and Ser283 variants (**Fig. 4A**). NP has six tryptophans that are relatively evenly distributed across the structure, and both NP variants exhibited a fluorescence emission maximum at 315 nm in the native state. Denaturation caused red-shifting of the emission maxima to 344 nm, and resulted in a decrease in emission intensity. The Pro283 and Ser283 NP fluorescence spectra were essentially indistinguishable both in the native and denatured conformations (**Fig. 4A**).

**Figure 4.**
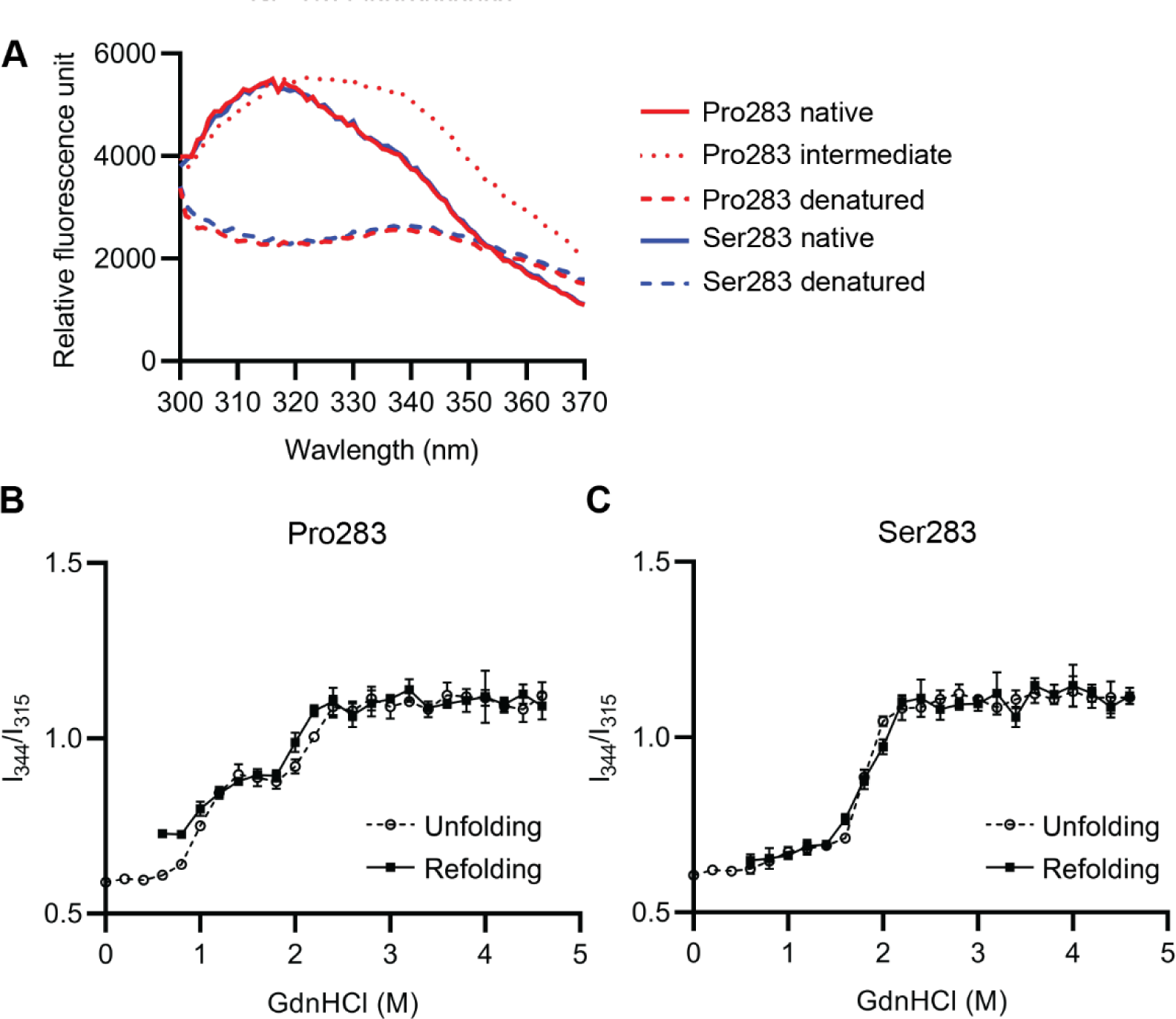
Pro283 NP folds in a three-state transition, while Ser283 NP folds in a two-state transition. (**A**) Intrinsic fluorescence spectra of native, intermediate (1.8 M GdnHCl) (in the case of Pro283 NP), and denatured NP. (**B**, **C**) I_344_/I_315_ plotted against GdnHCl concentrations for the unfolding and refolding transitions of Pro283 (**B**) and Ser283 (**C**) at room temperature.

To monitor unfolding and refolding transitions, we used the ratio of fluorescence intensity at 344 nm (I_344_) to fluorescence intensity at 315 nm (I_315_). Native protein fluorescence exhibited I_344_/I_315_ of ∼0.6, while denatured protein had I_344_/I_315_ of ∼1.1. The unfolding equilibrium was first measured by diluting native protein into refolding buffer with various GdnHCl concentrations at 25 °C, with buffer additives optimized to support maximal refolding (30% glycerol, 5 mM EDTA, and 5 mM dithiothreitol [DTT]). Strikingly, Pro283 exhibited three-state unfolding, while Ser283 exhibited two-state unfolding (**Fig. 4B** and **4C**). For Pro283, the first midpoint of transition occurred at 1.0 M GdnHCl and the second at 2.2 M GdnHCl (**Fig. 4B**). On the other hand, the intermediate conformer was not observed in the unfolding transition for Ser283, with a single transition midpoint at 1.8 M GdnHCl (**Fig. 4C**).

We also monitored the refolding transition by diluting NP denatured in 6 M GdnHCl into 0 M GdnHCl buffer (the dilution was performed at a 1:9 ratio, so the lowest tested concentration was 0.6 M). The refolding transition of Pro283 resembled the unfolding transition above 1 M GdnHCl with weak hysteresis, with the second transition midpoint at 2.0 M GdnHCl. (**Fig. 4B**). Below 1 M GdnHCl, Pro283 was unable to fully fold back into its native conformation, with I_344_/I_315_ at 0.6 M and 0.8 M GdnHCl around 0.7. In contrast, the refolding transition of Ser283 was almost identical to the unfolding transition at all GdnHCl concentrations tested with a single transition midpoint at 1.8 M (**Fig. 4C**).

### Pro283 NP is more aggregation-prone than Ser283 NP during unfolding and refolding

To characterize the intermediate conformers only observed with Pro283 NP, we examined the raw emission spectra. The intermediate species had an emission spectrum with a broad peak across 314 nm–338 nm, and increased fluorescence intensity compared to the native protein (**Fig. 4A**). Since tryptophan emission often increases with hydrophobic burial which leads to less collisional quenching, we suspected that the increased fluorescence emission of the intermediate species is indicative of aggregate formation (*27*).

To assess if Pro283 NP formed pelletable aggregates, we centrifuged the samples at 20,000 × *g* and compared the fluorescence of the supernatant to the fluorescence of the samples prior to centrifugation. First, during the equilibrium unfolding of NP, we observed that the fluorescence intensity decreased upon centrifugation for Pro283 NP between 0.6 M – 2.2 M GdnHCl, with the largest difference at 1.4 M GndHCl (**Fig. 5A**). This result indicates that the intermediate conformer formed pelletable aggregates. The aggregation was not mediated by intramolecular or intermolecular disulfides, as the buffer contained 5 mM DTT. On the other hand, for Ser283 NP the fluorescence spectra did not change upon centrifugation throughout all GdnHCl concentrations tested, indicating that Ser283 NP does not form pelletable aggregates during unfolding (**Fig. 5B**) at room temperature. Second, during the equilibrium refolding of NP, we also observed a decrease in the fluorescence intensity upon centrifugation with Pro283 NP (**fig. S6A**). Notably, a larger fraction of Pro283 formed pelletable aggregates during refolding than during unfolding. We also observed a small fraction of Ser283 NP forming pelletable aggregates during refolding (**fig. S6B**).

**Figure 5.**
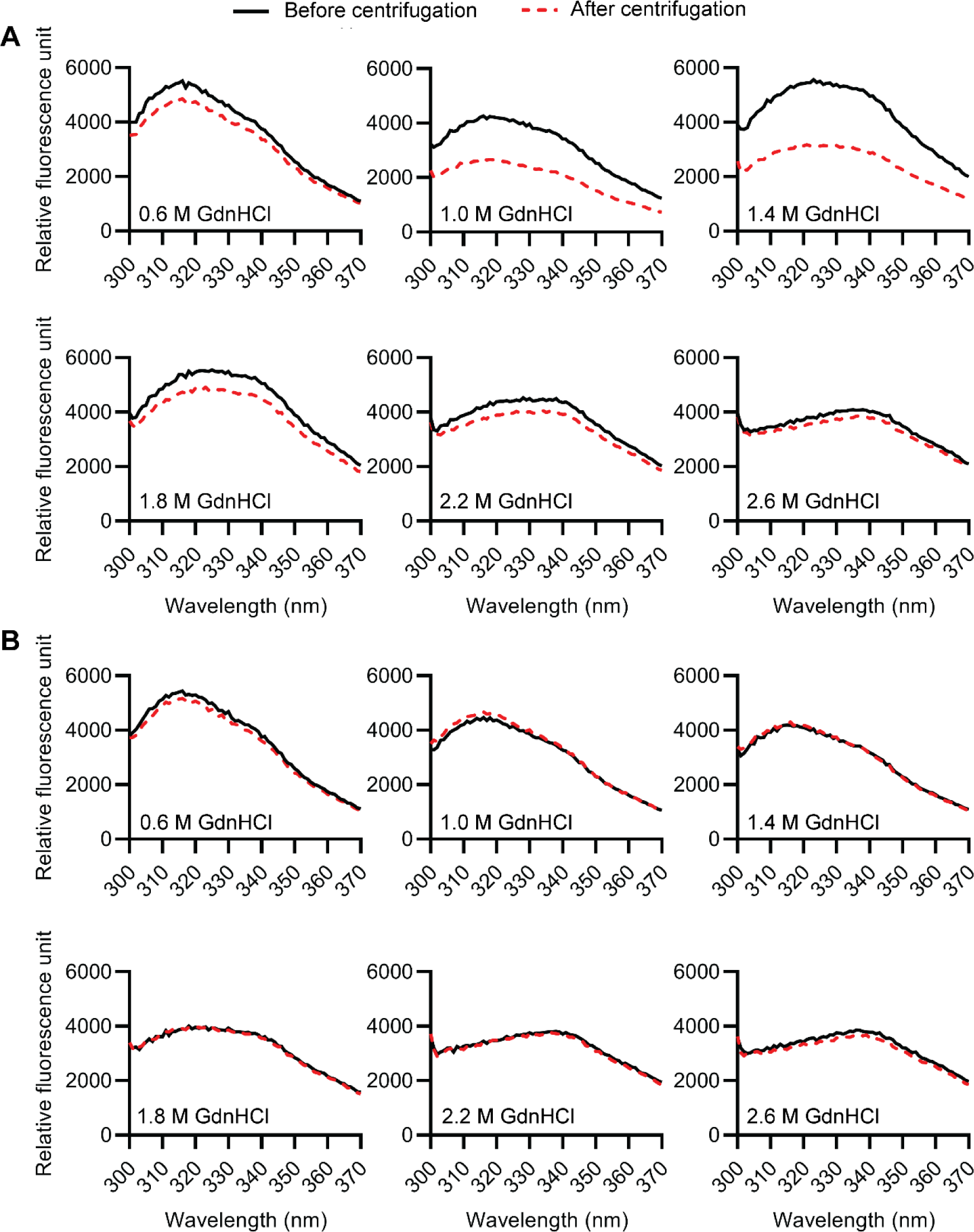
During unfolding at room temperature, Pro283 NP forms pelletable aggregates while Ser283 NP does not. Trp fluorescence spectra for (**A**) Pro283 and (**B**) Ser283 NP at various concentrations of GdnHCl during unfolding at room temperature, before (solid line) and after (dashed line) centrifugation to remove pelletable aggregates.

We next asked whether the higher aggregation propensity of Pro283 NP was reproducible at body temperature by repeating the GdnHCl-mediated denaturation at 37 °C. During unfolding at 37 °C, both variants exhibited three-state transitions (**fig. S7A** and **S7B**). Still, there were differences between Pro283 NP and Ser283 NP unfolding that indicated the Pro283 NP was more aggregation-prone. First, Pro283 NP had a lower first transition midpoint than Ser283 NP (0.3 M GdnHCl for Pro283 vs. 0.6 M GdnHCl for Ser283). This result indicates that a smaller degree of partial unfolding is sufficient for Pro283 NP to form an aggregation-prone intermediate compared to Ser283. Indeed, when we compared the fluorescence spectra before and after centrifugation, Pro283 NP fluorescence entirely disappeared at 0.4 M and 0.8 M GdnHCl, indicating that all of Pro283 NP formed pelletable aggregates at these conditions (**fig. S8A**). On the other hand, ∼85% of Ser283 NP remained soluble at 0.4 M GdnHCl, and ∼15% of Ser283 NP remained soluble at 0.8 M GdnHCl(**fig. S8B**). Second, while the difference was not substantial, Pro283 NP assumed the aggregation-prone intermediate conformation across a wider GdnHCl range than did Ser283 NP (0.3 M–1.8 M for Pro283 vs. 0.6 M–1.9 M for Ser283). In sum, Ser283 NP displayed a consistently lower propensity for aggregation under the same denaturation conditions.

Next, we monitored NP refolding at 37 °C. For Pro283 NP, the refolding transition was identical to the unfolding transition (**fig. S7A**). At the concentrations tested (final [GdnHCl] ≥ 0.6 M), Pro283 NP could not escape the intermediate conformation and spontaneously refold. This observation was confirmed by comparing the fluorescence emission before and after centrifugation, where the entire protein was lost due to aggregation at 0.8 M GdnHCl (**fig. S8C**). On the other hand, the Ser283 NP transition exhibited hysteresis when compared to the unfolding transition, indicating that the folding and unfolding pathways are likely to be different (**fig. S7B**). For example, at 0.8 M GdnHCl, only ∼15% of Ser283 NP remained soluble during unfolding (**fig. S8B**; 0.8 M GdnHCl), while ∼37% of Ser283 NP remained soluble during refolding (**fig. S8D**; 0.8 M GdnHCl).

Finally, since both Pro283 NP and Ser283 NP exhibited at least partial aggregation upon refolding at 37 °C, we tested if there was a difference in the rate of aggregation at 0.6 M GdnHCl. For this purpose, we measured solution turbidity change upon refolding by monitoring OD_350_, which has been used as a direct measure of static light scattering from formation of macromolecular aggregates (**fig. S9**) (*28–31*). Consistent with fluorescence data, we saw a higher maximum OD_350_ for Pro283 NP than for Ser283 NP. Furthermore, we observed that the lag time for aggregation was greater for Ser283 NP than for Pro283 NP. For Pro283 NP, we observed OD_350_ increase ∼3 mins after mixing, while for Ser283 NP we observed OD_350_ rising ∼12 mins after mixing. Next, we excluded the data points corresponding to the lag time, and fitted the remaining data to a single exponential equation. Interestingly, the rate constants were essentially identical for Pro283 NP (0.0429/min) and Ser283 NP (0.0421/min). Together, these data indicate that, during refolding, Pro283 NP requires less time to begin aggregating, but the rate of aggregation is similar for Pro283 NP and Ser283 NP once aggregation begins. Overall, our fluorescence spectra and turbidity measurements indicate that Pro283 NP is remarkably more aggregation-prone than Ser283 NP during unfolding and refolding regardless of temperature.

## Discussion

Pro283 is a substitution on influenza NP whose fitness is influenced by multiple important selection pressures. In the presence of the innate antiviral protein MxA, Pro283 enables NP to evade MxA (*7*). Simultaneously, influenza encoding Pro283 NP exhibits reduced fitness, and the viability of influenza encoding Pro283 NP is linked to both temperature and the availability of HSF1-regulated host chaperones (*7, 13, 15*). Thus, this work is, to our knowledge, the first to experimentally elucidate how a viral protein variant with a critical adaptive function exhibits biophysical defects that may be addressed by host chaperones.

### Pro283 substitution on influenza NP changes the folding pathway of NP without altering the native structure

Our results show that Pro283 NP exhibits a high propensity to aggregate during both unfolding and refolding, indicating that the substitution alters the folding pathway of NP by introducing an intermediate conformer that forms pelletable aggregates. The high aggregation propensity of Pro283 was observed both at room temperature and at 37 °C. At room temperature, Pro283 NP formed pelletable aggregates while Ser283 NP did not. At 37 °C, both variants were capable of forming pelletable aggregates, but Pro283 NP formed aggregates at a lower GdnHCl concentration when compared to Ser283 NP, as well as when compared to Pro283 NP at room temperature. This result indicates that at higher temperatures, a smaller degree of unfolding is sufficient to induce Pro283 NP to form terminal aggregates and render the protein nonfunctional. A single amino acid substitution can often dramatically alter the aggregation propensity of amyloidogenic proteins (*32–34*), but such cases have not been widely characterized for immune-escape variants of viral proteins.

Our biophysical experiments revealed several other important consequences of the Pro283 substitution for NP. First, Pro283 NP denatures and aggregates at a lower temperature than Ser283, both in test tubes and in cells (**Figure 2**), consistent with influenza encoding Pro283 exhibiting decreased fitness particularly at a febrile temperature (*13*). This observation also agrees with our unfolding/refolding data, as thermal energy can promote transient, partial unfolding, which in the case of Pro283 NP triggers aggregation. Second, our X-ray crystal structures and NMR spectra showed that Pro283 NP does not substantially disrupt the native structure of NP. This result may be expected for a single amino-acid substitution, but is surprising considering that site 283 is located at the center of an α-helix (**Figure 3**). It is possible that the packing of the other parts of the protein against the helix help to stabilize the α-helical conformation enough to allow accommodation of the normally helix-breaking proline residue (*25*). These potentially stabilizing interactions may not be present in a partially unfolded or a dynamic alternate conformation of the protein, leading to the higher aggregation propensity of Pro283 NP. Third, NMR spectra indicated that Pro283 NP and Ser283 NP display different dynamics in their native states, and with Ser283 NP exhibiting more motion in the μs–ms regime. While we do not have information about the specific residues involved in the motion, the difference in in-solution dynamics is consistent with the two variants of NP exhibiting different folding trajectories. Alternatively, it is possible that the timescale of motion most relevant to aggregation of NP is not in the μs–ms range, and such dynamics could not be captured in our NMR experiments despite being more prevalent in Pro283.

Notably, the change in aggregation propensity introduced by Pro283 may have a more severe consequence for influenza viability during the viral replication cycle *in vivo*. The recombinantly expressed NP in our work contained modifications to prevent RNA binding and promote monomeric expression (*13, 17, 35*). On the other hand, in cells, NP undergoes oligomerization mediated by a flexible tail loop, which is critical for viral ribonucleoprotein complex assembly (*3, 17*). The absence of modifications that favor monomeric expression, together with the high local concentration of NP, could further enhance the aggregation propensity of Pro283 NP *in vivo*.

### How host chaperones potentially address Pro283 NP-associated biophysical defects

Emerging studies since 2010s began to show that host proteostasis networks can shape the evolution of viral proteins (*13, 36–41*). Nevertheless, while it is evident that host proteostasis factors, particularly chaperones, can define the fitness of viral protein variants, the molecular mechanism has been largely left unclear (*16*). Our biophysical characterization of Pro283 NP suggests that host-cell chaperones can potentially rescue Pro283 NP by stabilizing an aggregation-prone conformation, or promoting the rapid conversion of the aggregation-prone conformation to a folded conformation In addition, it is important to note that the unfolding/refolding studies performed in this work are *in vitro*, and aggregation-prone intermediates formed during *in vitro* refolding sometimes are not observed when protein are allowed to co-translationally fold (*42*). It is possible that co-translational binding of chaperones can prevent the aggregation-prone intermediate from forming in the first place (*43*).

Our work does not yet identify the specific host chaperones that rescue the biophysical defects of Pro283 NP. Since HSF1 regulates the expression of a broad change of proteostasis factors (*44–46*), potential candidates include heat shock proteins that can bind to the partially folded conformation and prevent aggregation and folding enzymes like peptidyl prolyl isomerases that catalyze this often rate-limiting step folding. Indeed, it has been observed that the nucleoprotein of SARS Cov-2 virus heavily interacts with peptidyl prolyl isomerases like CypA and PIN, although it is not yet clearly known whether these enzymes play an important role in the nucleoprotein folding (*47*). The chaperones that can rescue Pro283 NP may have homologous proteins in *E. coli*, since Pro283 NP could be recombinantly expressed in *E. coli* without a noticeable yield decrease.

### Implications of the biophysical cost of the Pro283 substitution for immune escape of NP

The high-level phenotypic consequence of a mutation (e.g., fitness) is a cumulative outcome of the mutation’s effect on multiple component phenotypes (e.g., MxA escape, thermal stability, aggregation propensity, etc) (*48*). Additional selection pressures, such as treatment with nucleozin, which is an antiviral drug that induces nucleoprotein aggregation (*49*), can further convolute how the Pro283 substitution affects the fitness of the influenza. It will be also interesting to investigate what, if any, evolutionary strategies NP can use to address Pro283-induced folding defects when host chaperone assistance is not available, such as stabilizing epistatic mutations.

Furthermore, both our data and another crystal structure of H3N2 NP (*26*) show that the three MxA-resistant substitutions identified in the 1918 influenza pandemic (*7*) do not perturb the native structure of NP. This result strongly suggests that MxA resistance via Pro283 substitution is not mediated by secondary structure disruption, but rather by atomic interactions on the individual amino-acid level. A similar case has been observed for HIV-1 Vif, where the residues lining the surface of an α-helix were shown to be critical for APOBEC3G degradation and viral infectivity, owing to a combination of residue-specific aliphatic interactions, aromatic base stacking, and hydrogen bonds (*50*). So far, only an unpublished computational study predicted that Pro283 substitution would weaken the aliphatic interaction with F561 of MxA (*51*). Structures of NP-MxA complex in the presence and absence of key MxA-escape mutations would greatly advance our understanding of the antiviral mechanism of MxA and how NP evolved to evade MxA.

## Materials and Methods

### Plasmids

Wild-type influenza NP (Pro283) was expressed from a pET28b(+) expression vector encoding monomeric A/Aichi/2/1968 NP as previously described (*13*). The Pro283Ser amino substitution was introduced to this vector via the QuikChange II XL Site-Directed Mutagenesis Kit (Agilent) as previously described (*13*). The nucleotide sequence of the origin of replication with the mutation highlighted is provided in **Supplementary File 1**. The NP construct included a C-terminal 6×-His tag for Ni-affinity purification, and R416A substitution and deletion of residues 2–7 for RNA-free, monomeric expression. HDM_FLAG_Aichi68_NP_IRES_mCherry NP plasmid used for construction of NP-encoding adenovirus was a kind gift from Professor Jesse D. Bloom (Fred Hutchinson Cancer Research Center, Seattle WA U.S.A.).

### Cell Lines and Reagents

The following cell lines were only used for cellular thermal shift assays: MDCK (American Type Culture Collection) and HEK 293A (Thermo Fisher) cells were cultured at 37 °C in a 5% CO_2_(g) atmosphere in complete DMEM media (Corning) supplemented with 10% fetal bovine serum (CellGro), 1% penicillin/streptomycin (CellGro), and 2 mM L-glutamine (CellGro). The MDCK cells used in this work had the identical cellular background to the MDCK cells used in the deep mutational scanning work that first identified that the relative fitness of influenza Pro283 NP depends on temperature and host proteostasis environments (*13*).

### Recombinant Expression of NP

For recombinant expression in *E. coli*, BL21(DE3) chemically competent cells were transformed with 1 μL of purified plasmid and incubated overnight on LB-kanamycin agar plates. Colonies were used to inoculate 50 mL LB-kanamycin cultures overnight. Then, 10 mL starter cultures were used to inoculate 1 L LB-kanamycin cultures, which were shaken at 37 °C until an OD_600_ of 0.6–0.8 was attained. Cultures were then induced with 500 μM IPTG (Sigma) overnight at 20 °C. Cells were pelleted by centrifuging at 8,000 × g for 10 minutes at 4 °C, Dounce homogenized, and lysed by sonication in 50 mL of lysis buffer (50 mM Tris at pH 7.5, 300 mL NaCl, 5 mM imidazole, 10% glycerol, 0.2 mM 2-mercaptoethanol, 1 mM PMSF, and Pierce Protease Inhibitor Tablets [Thermo Fisher]). Cells were sonicated for 5 min (30% amplitude, 10 s on, 10 s off; Branson Digital Sonifier). Lysates were then clarified for 30 min by centrifuging at 10,000 × g at 4 °C, and incubated with 50 μL of DNase I (1 unit/μL, Thermo Fisher Scientific) and 50 μL of RNase A (10 mg/mL, Thermo Fisher Scientific) for 20 minutes at rt. The lysates were then incubated on Ni-NTA (Millipore) columns for 60 min at 4 °C and washed with lysis buffer. Proteins were eluted using elution buffer (50 mM Tris at pH 7.5, 300 mL NaCl, 250 mM imidazole, and 10% glycerol). Elutions were dialyzed overnight (against 50 mM Tris at pH 7.5, 300 mL NaCl, and 10% glycerol) at 4 °C to remove imidazole, and then purified over a heparin column (Cytiva HiPrep Heparin FF 16/10). NP eluted approximately with 10% Buffer A (50 mM Tris at pH 7.5, 1 mM EDTA, 10% glycerol, 5 mM 2-mercaptoethanol) and 90% buffer B (50 mM Tris at pH 7.5,1 M NaCl, 1 mM EDTA, 10% glycerol, 5 mM 2-mercaptoethanol). Elutions were further purified by size exclusion chromatography (Bio-Rad Enrich SEC 650). Protein purity was checked by 4%/12% SDS-PAGE. Concentrations were determined by measuring A280 with a BioTeK Synerge H1 microplate reader and BioTek Take3 microvolume plate (Agilent) using the calculated extinction coefficient of 55,350 M^−1^cm^−1^. Purified proteins were concentrated using Amicon Ultra 3K MWCO filters (Millipore) in storage buffer (50 mM Tris-HCl pH 7.6, 50 mM NaCl, 1 mM DTT) and stored at −80 °C.

For recombinant expression of ^2^H^13^C^15^N-labeled NP, we used a modified version of a previously developed high-cell-density expression method (*52*). 2.5 mL of overnight starter cultures were used to inoculate 250 L LB-kanamycin cultures, which were shaken at 37 °C until an OD_600_ of –1.6 was attained. The cultures were centrifuged and the pellets were washed with D_2_O. The pellets were then resuspended in 250 mL M9 media in D_2_O (50 mM Na_2_HPO_4_, 25 mM KH_2_PO_4_, 10 mM NaCl, 5 mM MgSO_4_, 0.2 mM CaCl_2_, 0.1% w/v ^15^NH_4_Cl. 1% w/v U-^13^C-U-^2^H-D-glucose. 0.1% v/v trace metal solution [50 mM H_3_BO_4_, 0.2 mM CoCl_2_, 1 mM CuSO_4_, 1 mM MnCl_2_, 1 μM Na_2_MoO_4_, 2 mM ZnCl_2_, and 1 mM FeSO_4_], and 1% v/v MEM vitamin solution [Thermo Fisher, lyophilized and dissolved in D2O]). The culture was grown at 37 °C for 1 h and was induced with 500 μM IPTG for 24 h at 20 °C. The harvest, lysis, and purification steps were identical to the unlabeled NP expression.

### Engineering NP-encoding Adenovirus

The described adenoviruses were only used for ectopic expression of NP for cellular thermal shift assays, as MDCK cells are not transfectable. Adenoviral destination vectors encoding C-terminal HA-tagged Pro283 NP, C-terminal HA-tagged Ser283 NP, and untagged Pro283 NP were used to produce adenovirus according to the manufacturer’s instructions (Life Technologies ViraPower Adenoviral Expression System). NP-encoding sequence was cloned from plasmid HDM_FLAG_Aichi68_NP_IRES_mCherry.

Viruses were amplified in HEK293A cells, and were titered using a plaque assay (described below) and used for subsequent experiments. For adenovirus titering by plaque assay, HEK 293A cells were seeded at a density of 1 × 10^6^ cells/well in a 6-well plate 24 h prior to infection with a serial 10-fold dilution of passage 2 or 3 adenoviruses. Media was removed 6 h post-infection and the cells were overlaid with 0.4% agarose in DMEM. When plaques were observed (typically after 7–8 d), the cells were stained with 200 μL of 5 mg/mL solution of 3-(4,5-dimethylthiazol-2-yl)-2,5-diphenyltetrazolium bromide (MTT) stain (Research Products International) in PBS. After 16 h the plaques were counted manually and viral titer was calculated in plaque forming units per μL viral stock (*53*).

### Cellular Thermal Shift Assay

MDCK cells at 2 × 10^6^ cells/plate were transduced with adenoviruses to produce Aichi/1968 (H3N2) influenza A NP. Media was changed 16 h post-transduction. 24 h later, cells were trypsinized and harvested, pelleted (1000 × g for 5 min), and resuspended in 5 mL of PBS. 100 μL samples of cells at 3 × 10^6^ cells/mL concentration were heated in a thermocycler for 3 min, then cooled to RT for 3 min. Samples were promptly lysed using five freeze-thaw cycles, then centrifuged (20,000 × g for 20 min). The resulting supernatant was separated on a reducing SDS-PAGE gel. Protein bands were transferred to a nitrocellulose membrane and immunoblots were probed with anti-HA primary antibody (Thermo Fisher 26183,1:10000) and anti-mouse secondary antibody (Abcam ab205719, 1:1000), imaged with a ChemiDoc (Bio-Rad), and quantified using Image Studio.

### Dynamic Light Scattering

A DynaPro Nanostar (Wyatt Technology) was used for dynamic light scattering measurements. The laser was set at λ = 658 nm with the detector angle at 90° for scattering measurements. The samples were centrifuged at 21,000 × g for 15 minutes prior to measurement to remove dust and large-particle contaminants. The spectra were acquired in a disposable polystyrene cuvette at room temperature at a protein concentration of 1.5 mg/mL in 50 mM Tris at pH 7.5, 50 mM NaCl, 1 mM DTT buffer (Acquisition time = 5 s, number acquisition = 20).

### Analytical Ultracentrifugation

Sedimentation-velocity analytical ultracentrifugation experiments were performed at 20 °C with an XL-I analytical ultracentrifuge (Beckman-Coulter) and an An50 Ti rotor. NP was dissolved in 50 mM Tris at pH 7.5, 50 mM NaCl, and 1 mM DTT at a final protein concentration of 0.8 mg/mL. Cells with 12-mm charcoal-filled Epon double-sector centerpieces were used with 420 μL buffer and 410 μL NP. Sedimentation boundaries were measured at a speed of 40,000 rpm by scanning absorbance at 280 nm between 5.8 and 7.2 cm radial distances with a step size of 0.002 cm. Sedimentation coefficient distributions of Lamm equation solutions were generated using Sedfit (v. 16.5) using a continuous *c*(s) distribution (*54*). The following parameters were used for fitting: *S*_min_ = 0, *S*_max_ = 10, buffer density = 1.005, relative buffer viscosity = 1.0108, best fit friction ratio = 1.263 (Pro283) or 1.200 (Ser283). The artificial peak at *S* < 0.1 was excluded when calculating the total percentage of protein.

### Differential Scanning Fluorimetry

Differential scanning fluorimetry was performed using a Light Cycler 480 II Real-time PCR System (Roche). NP (in 50 mM Tris at pH 7.5, 50 mM NaCl, 1 mM DTT) was mixed with 5000× SYPRO Orange stock solution (Thermo Fisher), at a final volume of 10 μL and a final concentration of 50× SYPRO Orange and 0.05–1 mg/mL NP. An excitation wavelength of 465 nm and an emission wavelength of 580 nm were used. Thermal shift assays were performed at a ramp rate of 0.05 °C/sec from 37–60 °C in continuous acquisition with 10 acquisitions per °C. Assays were performed in 384-well plates in five technical replicates.

### Crystallography

NP crystals were grown in hanging drops under the condition specified in **Table 1**. 1 μL of NP (13.5 mg/mL in 50 mM Tris-HCl at pH 7.5, 50 mM NaCl, 1 mM DTT) was mixed with 1 μL of reservoir solution, and rod/star shaped crystals appeared in 3 d. Crystals were frozen in liquid nitrogen directly from the drop without cryo-protection. Diffraction data were collected at 100 K crystals at the Advanced Photon Source at Argonne National Laboratory, NE-CAT beamline 24-IDE. Data were indexed, integrated, and scaled to 2.90 Å (Pro283) or 3.09 Å (Ser283) with XDS (*55*). Structures were solved by molecular replacement using the Phenix implementation of Phaser (*56, 57*) with WSN monomeric NP (PDBID: 3ZDP) as the search model. The structure was refined with iterative rounds of refinement in Phenix (*57*), model building with Coot (*58*), and validated by MolProbity (*59*). **Table 2** reports data and refinement statistics. The structures are deposited in the PDB with the identifiers 8TWP (Pro283) and 8TWR (Ser283). RMSD values, inter-atom distances, and dihedral angles were all obtained using PyMOL 2.3.3 (the PyMOL Molecular Graphics System Version 2.3.3 Schrodinger, LLC), and structure figures were also made with PyMOL 2.3.3. RMSD per residue values were calculated using the rmsdCA.py script (Y. Tong, rmsdCA, (2021), GitHub repository, https://github.com/tongalumina/rmsdca)

### NMR spectroscopy

A buffer in 90% H_2_O and 10% D_2_O consisting of 50 mM Tris, pH 7.5, 50 mM NaCl, and 1 mM DTT was used for all NMR samples. The 1D ^1^H NMR stability tests were performed with samples containing ∼190 μM NP. Samples were monitored via NMR over the course of 4–5 d. For perdeuterated nucleoprotein samples, labile deuterons of the protein were exchanged back to protons during the purification process. NMR data were collected at 25 °C with 300 μL samples containing ∼250 μM of either ^2^H-, ^13^C-, and ^15^N-uniformly labeled Pro283 NP or ^2^H-, ^13^C-, and ^15^N-uniformly labeled Ser283 NP. Experiments were performed using Shigemi tubes matched with D_2_O.

1D experiments were performed at either 500 MHz or 600 MHz, while 2D and 3D experiments were performed on an 800 MHz Avance Neo Bruker NMR spectrometer equipped with a TXO cryo-probe optimized for ^15^N detection. Chemical shifts were referenced relative to the spectrometer frequency, with the water resonance at ∼4.7 ppm. Pulses were calibrated using the standard Topspin protocol. For 1D ^1^H NMR experiments, the following parameters were used for each NMR spectrometer frequency: **1)** 500 MHz NMR spectrometer using the pulse program zgesgp (*60*): 15.61 ppm spectral width, frequency offset of 4.700 ppm, 32768 points, 64 scans, and DQD acquisition mode with a relaxation delay of 1.5 s. **2)** 600 MHz NMR spectrometer using the pulse program p3919gp (*61, 62*): 20.03 ppm spectral width, frequency offset of 4.691 ppm, 8192 points, 32 scans, and DQD acquisition mode with a relaxation delay of 1.5 s. **3)** 800 MHz NMR spectrometer using the pulse program zgesgp (*60*): 13.58 ppm spectral width, frequency offset of 4.728 ppm, 4096 points, 32 scans, and DQD acquisition mode with a relaxation delay of 1.5 s.

D ^1^H-^15^N TROSY (*63*) experiments were performed with the pulse program trosyetfpf3gpsi (*64–69*) with 16 scans and a relaxation delay of 1.2 s. The following parameters were set for the ^1^H dimension: 14.20 ppm spectral width, frequency offset of 4.720 ppm, 2048 points and DQD acquisition mode. The following parameters were used for the ^15^N dimension: 40.00 ppm spectral width, frequency offset of 118 ppm, 256 points and Echo-Antiecho acquisition mode.

1D ^1^H NMR data were processed and analyzed using MestReNova © (version: 14.2.1-27684). Multidimensional NMR data were processed using NMRPipe(*70*) and analyzed in NMRFAM-SPARKY (*71*) (Sparky).

### Equilibrium unfolding/refolding experiments

For equilibrium unfolding experiments, purified NP was diluted into 50 mM Tris at pH 7.5, 50 mM NaCl, 30% glycerol, 5 mM EDTA, and 5 mM DTT buffer containing 0–5 M GdnHCl. For equilibrium refolding experiments, purified NP was denatured by dialyzing against 6 M GdnHCl solution with 20 mM phosphate at pH 7.5. The denatured NP was then diluted into the aforementioned buffer. For both unfolding and refolding experiments, the protein-to-buffer ratio for dilution was 1:9 to give a final protein concentration of 0.25 mg/mL. The samples were incubated at room temperature for at least 5 h to ensure equilibrium had been reached.

Fluorescence emission spectra were recorded using a BioTeK Synerge H1 microplate reader and BioTek Take3 microvolume plate (Agilent) with an excitation wavelength of 274 nm. Turbidity was monitored using a BioTeK Synergy H1 microplate reader and a polyproline 96-well plates (Corning) with 100 μL sample volume and no agitation applied. Experiments were performed in technical quadruplicates.

### Statistical analysis

Unless indicated otherwise, experiments were performed in technical triplicates with replicates defined as independent experimental measurements from same biological samples. For differential scanning fluorimetry, *T*_agg_ values were obtained by performing a nonlinear regression to a Boltzmann sigmoid function using GraphPad Prism. The mean of *T_agg_* values was tested for significance using a 2-sample *t* test in GraphPad Prism. For cellular thermal shift assay, densitometric analysis of immunoblots were tested for statistical significance using a 2-sample *t* test in GraphPhad Prism. For equilibrium refolding and unfolding experiments, the I_344_/I_315_ vs GdnHCl concentration data were fit to a Boltzmann sigmoid curve (two-state transition) or a double Boltzmann sigmoid curve (three-state transition) using GraphPad Prism. GraphPhad Prism version 9.4.1 was used for all statistical analyses involving GraphPad Prism.

## Acknowledgments

This work is based upon research conducted at the Northeastern Collaborative Access Team beamlines, which are funded by the National Institute of a General Medical Sciences from the National Institute of Health (P30 GM124165). The Eiger 16 M detector on the 24-ID-E beam line is funded by a NIH-ORIP HEI grant (S10OD021527). This research used resources of the Advanced Photon Source, a U.S. Department of Energy (DOE) Office of Science User Facility operated for the DOE Office of Science by Argonne National Laboratory under Contract No. DE-AC02-06CH11357. The authors thank Dr. Angela Phillips for providing helpful feedback on the conception of this work, and Dr. Xuemei Huang for assistance with NMR experiments.

## Funding

Kwanjeong Overseas Graduate Fellowship (JY)

Massachusetts Institute of Technology Mathworks Fellowship (JY)

National Cancer Institute (Koch Institute Support (core) Grant P30-CA14051) (MDS)

National Institute of Environmental Health Sciences (MIT Center for Environmental Health Sciences (core) Grant P30-ES002109) (MDS)

National Science Foundation (CAREER Award 1652390) (MDS) National Institute of Health (R01 AI168166) (MDS)

National Institute of Health (R35 GM138382) (GTD, CH)

## Author contributions

Conceptualization: JY, YMZ, AMP, MDS

Methodology: JY, YMZ, CH, RAG, AMP, BEA, GTD, MDS

Validation: JY, YMZ, CH, RAG Formal analysis: JY, YMZ, CH, RAG

Investigation: JY, YMZ, CH, RAG, AMP, BEA

Resources: JY, YMZ, CH, RAG, AMP, BEA, GTD, MDS

Visualization: JY, YMZ, CH, RAG

Supervision: GTD, MDS

Project administration: GTD, MDS

Funding acquisition: GTD, MDS

Writing—original draft: JY, YMZ, GTD, MDS

Writing—review & editing: JY, YMZ, CH, RAG, AMP, GTD, MDS

## Data and materials availability

Atomic coordinates and structure factors of the NP crystal structures have been deposited in the Protein Data Bank under accession code 8TWP (Pro283) and 8TWR (Ser283). All data needed to evaluate the conclusions in this paper are available in the main text or the supplementary materials.

