## Supplemental Information for "The Immune-Evasive Proline 283 Substitution in Influenza Nucleoprotein Increases Aggregation Propensity Without Altering the Native Structure"

Jimin Yoon *et al.*

**This PDF file includes:**

Figs. S1 to S9

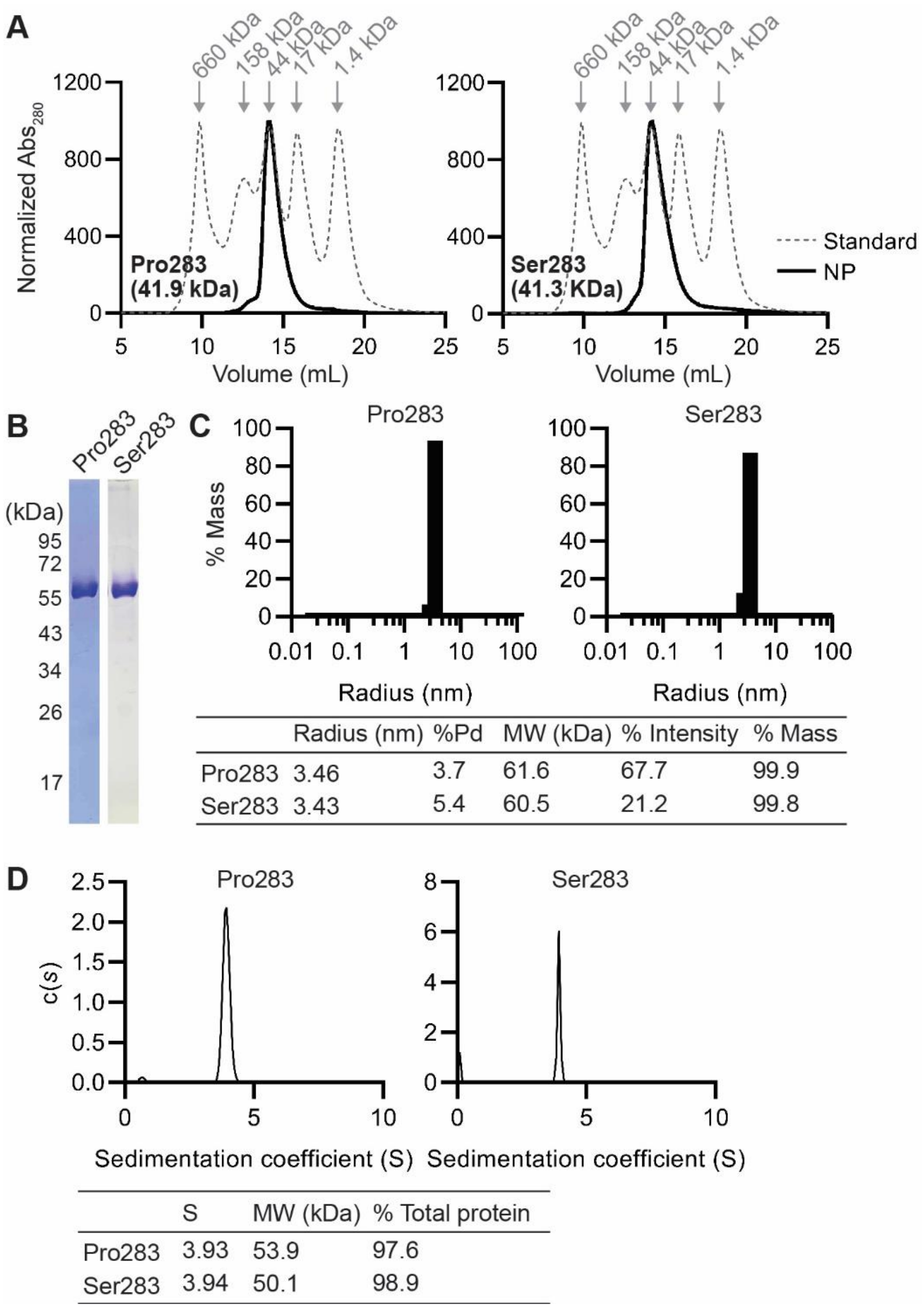


| | Radius (nm) | %Pd | MW (kDa) | % Intensity | % Mass |
|---|---|---|---|---|---|
| Pro283 | 3.46 | 3.7 | 61.6 | 67.7 | 99.9 |
| Ser283 | 3.43 | 5.4 | 60.5 | 21.2 | 99.8 |

| | S | MW (kDa) | % Total protein |
|---|---|---|---|
| Pro283 | 3.93 | 53.9 | 97.6 |
| Ser283 | 3.94 | 50.1 | 98.9 |

**Fig. S1. NP is recombinantly expressed and isolated as pure monomers.**

(**A**) Size exclusion chromatograms of purified Pro283 and Ser283. Grey dashed chromatograms are protein standards used for molecular weight prediction. (**B**) Coomassie-stained SDS-PAGE gel of size exclusion chromatogram fraction corresponding to NP. The single band corresponds to monomeric NP. (**C**) Dynamic light scattering data showing that both Pro283 NP and Ser283 NP are monodisperse with 3.7% and 5.4% polydispersity, respectively. (**D**) Sedimentation coefficient distribution generated from sedimentation velocity experiments showing a single peak at 3.9 S for both Pro283 and Ser283 NP, which constituted >97% of total protein.

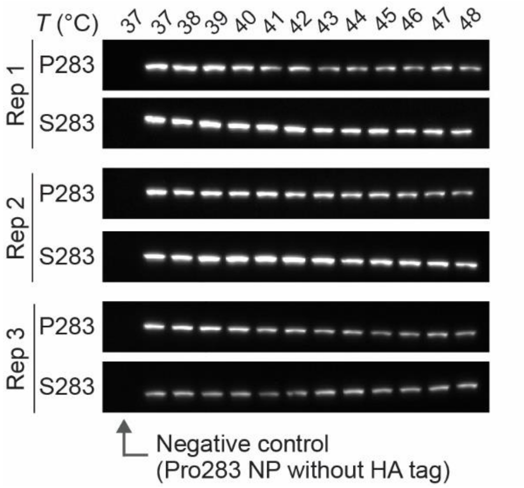


**Fig. S2. Cellular thermal shift assay immunoblots of Pro283 and Ser283 NP.**

Immunoblots from three technical replicates of NP cellular thermal shift assay. The first lane of each immunoblots corresponds to Pro283 NP without an HA-tag, which serves as the negative control for anti-HA antibody binding.

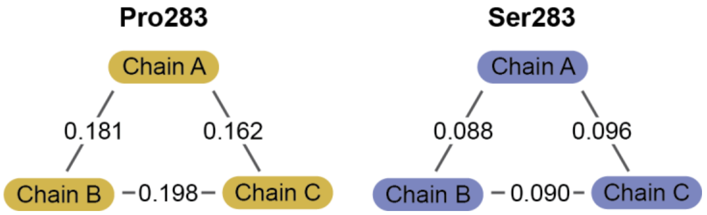


**Fig. S3. The molecules within an asymmetric unit are highly similar to each other for both Pro283 and Ser283 NP.**

RMSD values between chains in the asymmetric unit for the Pro283 and Ser283 NP crystal structures.

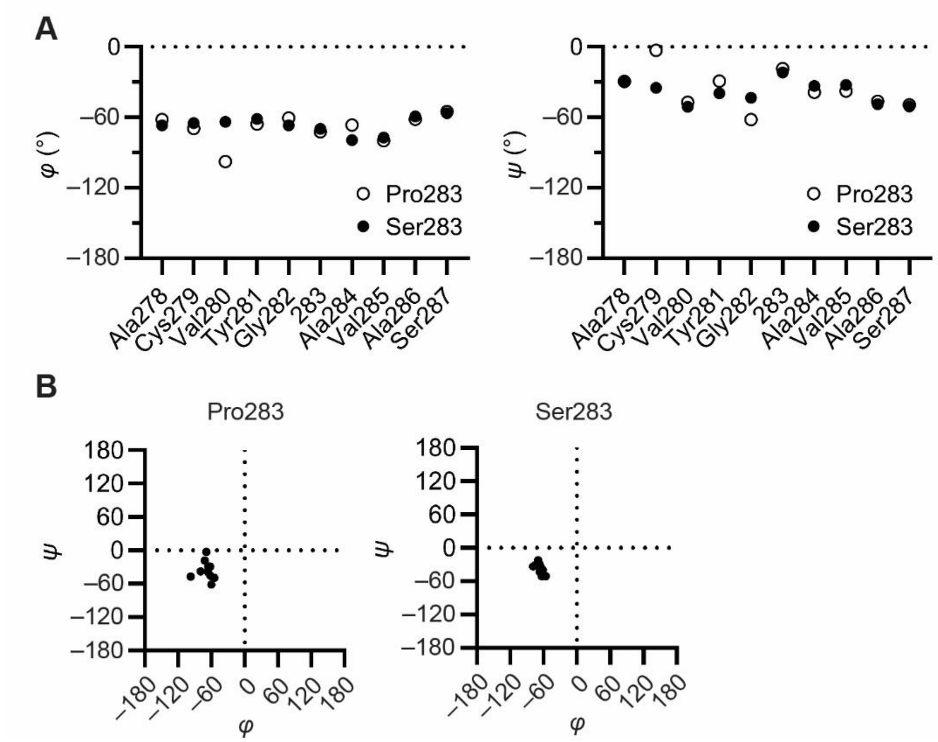


**Fig. S4. Pro283 substitution only slightly changes the dihedral angles of the α-helix containing site 283 (Ala278–Ser287).**

(**A**) *φ* and *ψ* angles of each residue for the α-helix containing site 283 (Ala278–Ser287). (**B**) Ramachandran plot of the α-helix containing site 283 (spanning Ala278 to Ser287).

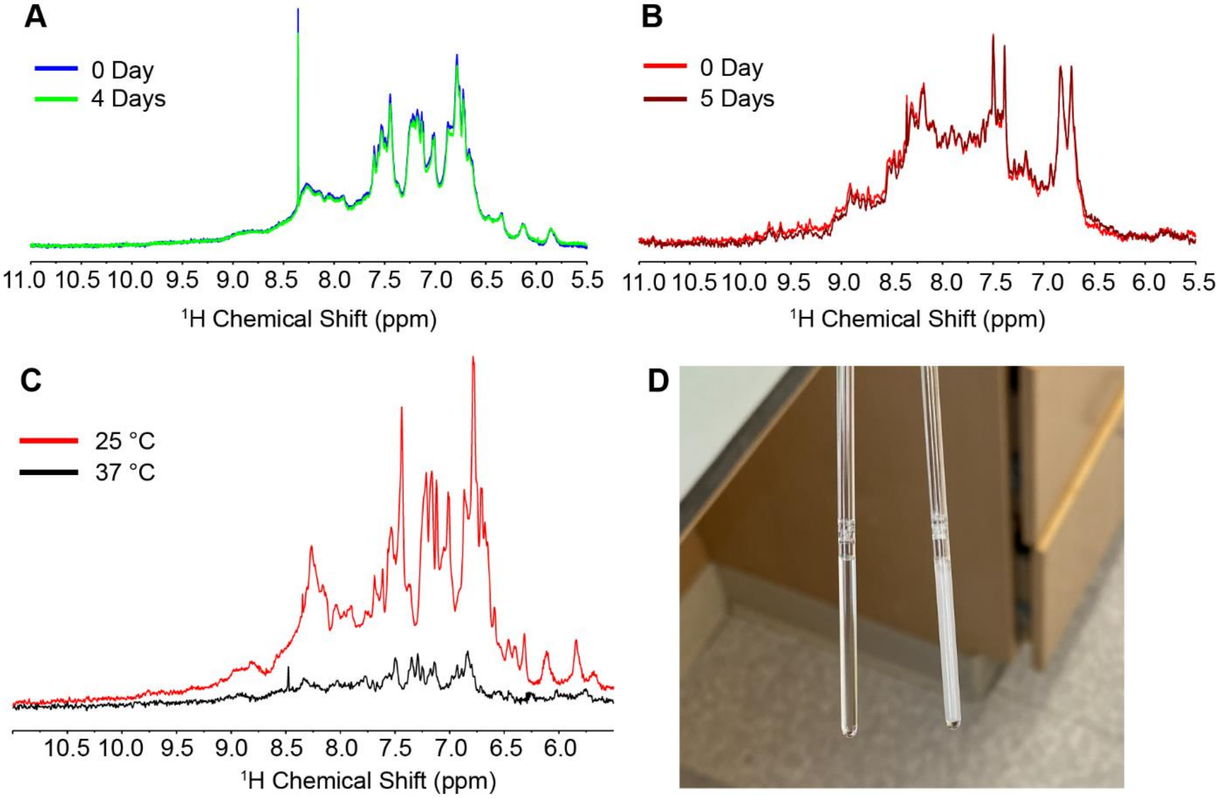


**Fig. S5. Stability of NP in buffer and concentration conditions suitable for NMR experiments at 25 °C and 37 °C.**

$^{1}$H-NMR of (**A**) Ser283 NP and (**B**) Pro283 NP over the course of several days at 298 K, demonstrating the stability of the samples under NMR conditions. (**C**) $^{1}$H-NMR of Pro283 NP at different temperatures. (**D**) Images of the NMR samples after experiments at 25 °C (clear sample) and at 37 °C (cloudy sample).

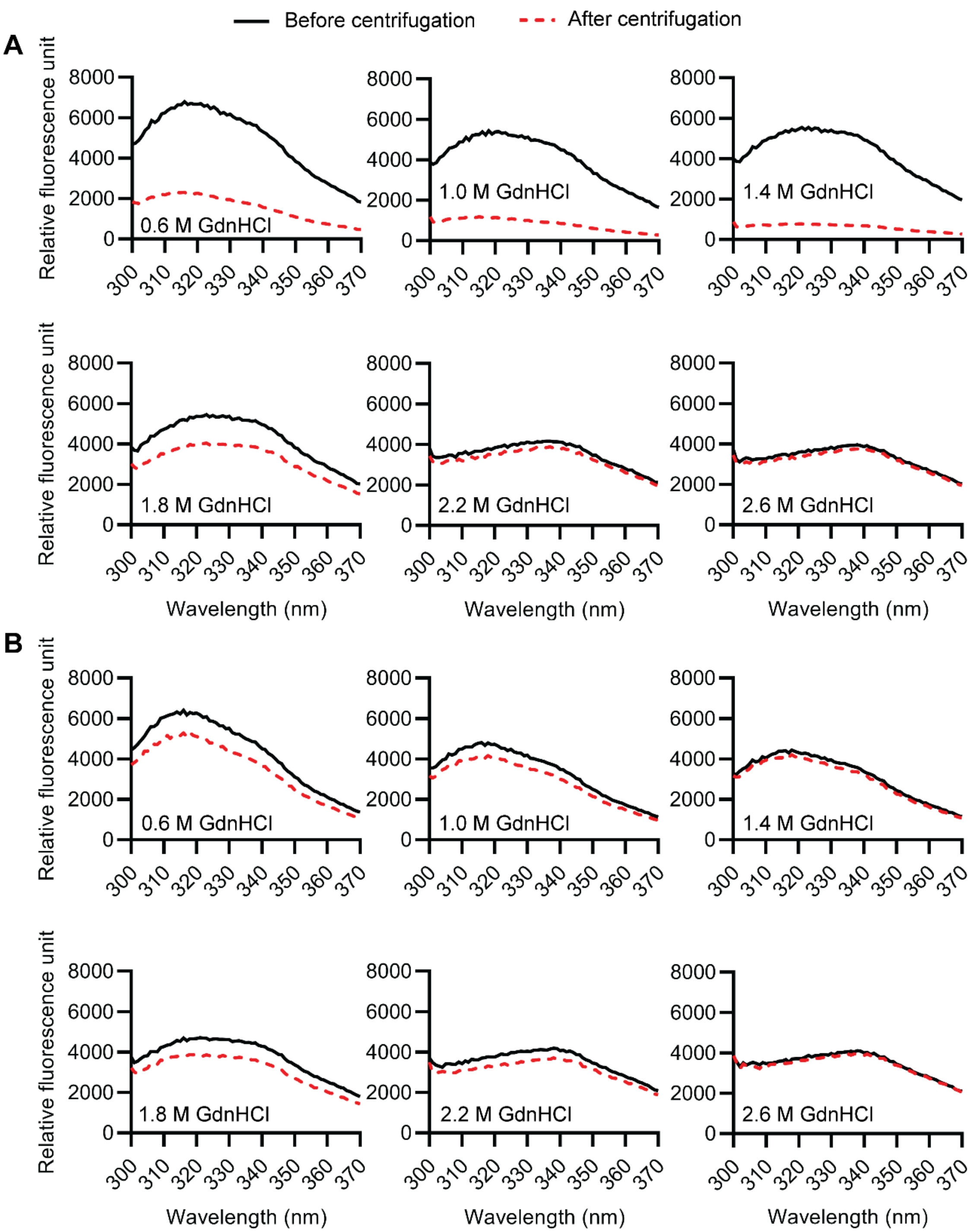


**Fig. S6. During refolding at room temperature, Pro283 NP forms pelletable aggregates substantially more than Ser283 NP.**

Trp fluorescence spectra for (**A**) Pro283 and Ser283 (**B**) NP at various concentrations of GdnHCl during refolding, at room temperature, before (solid line) and after (dashed line) centrifugation to remove pelletable aggregates.

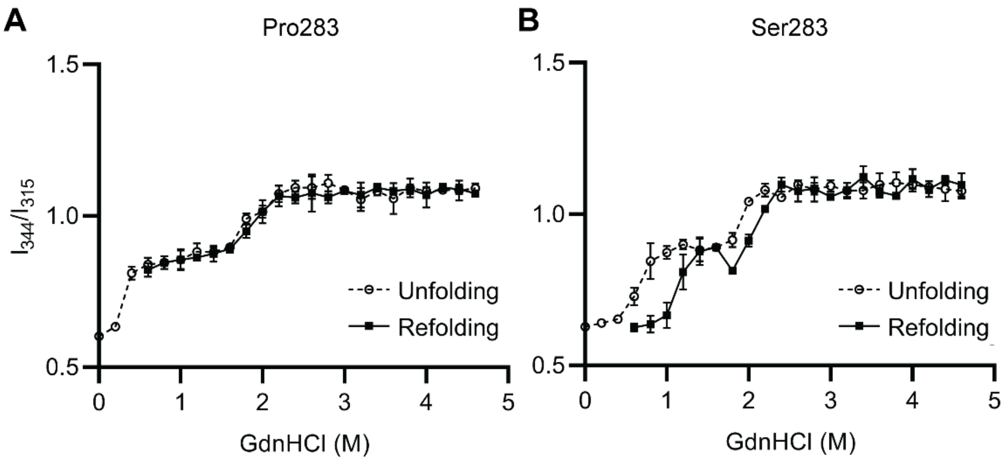


**Fig. S7. Pro283 NP is more aggregation-prone than Ser283 at 37 °C.**

$I_{344}/I_{315}$ plotted against GdnHCl concentrations for the unfolding and refolding transitions of Pro283 (**A**) and Ser283 (**B**) at 37 °C.

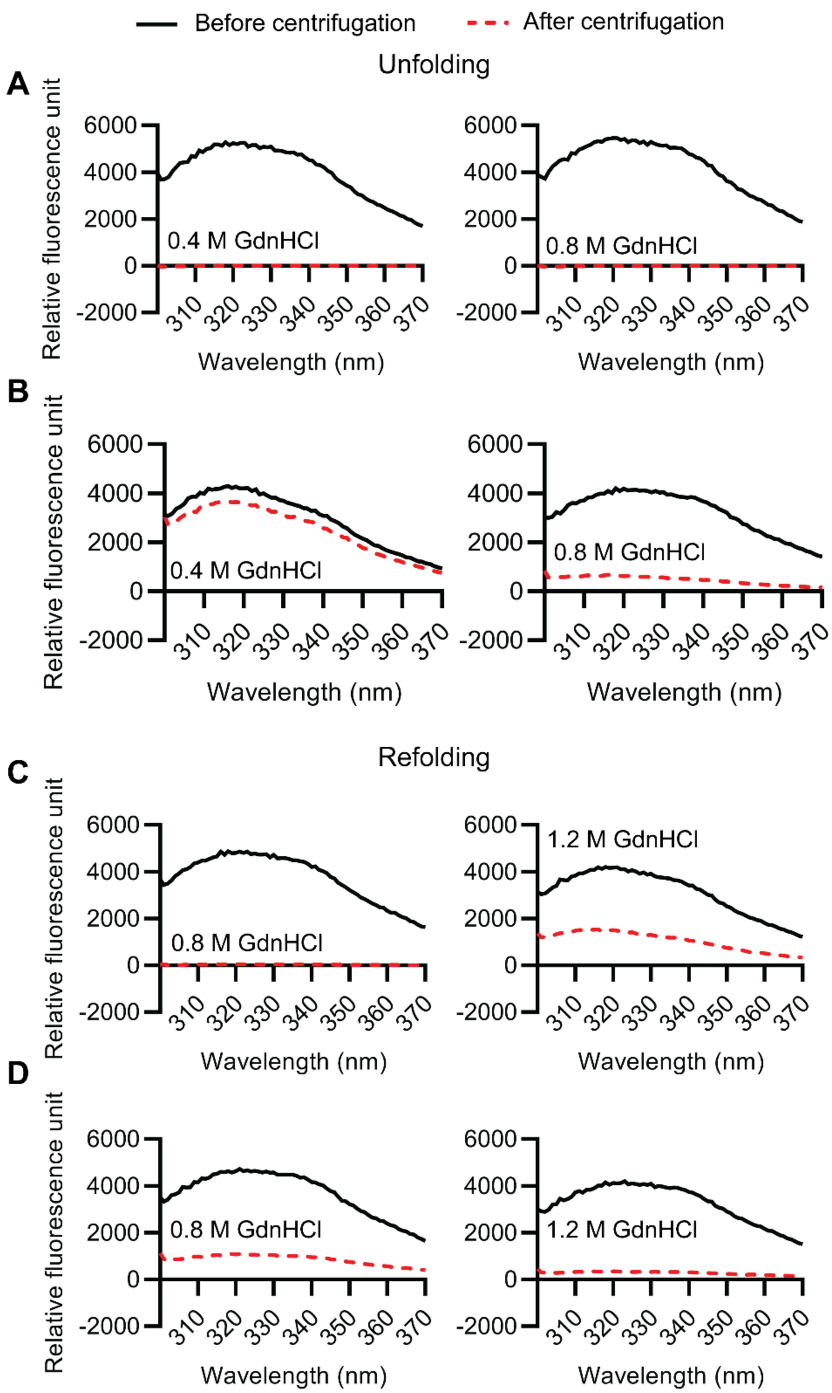


**Fig. S8. During unfolding and refolding at 37 °C, Pro283 NP forms pelletable aggregates at a lower GdnHCl concentrations than Ser283 NP.**

Trp fluorescence spectra during unfolding for (**A**) Pro283 and (**B**) Ser283 NP at various concentrations of GdnHCl, at 37 °C. Trp fluorescence spectra during refolding for (**C**) Pro283 and (**D**) Ser283 NP at various concentrations of GdnHCl, at 37 °C. Solid line and dashed line represent fluorescence spectra before and after centrifugation to remove pelletable aggregates, respectively.

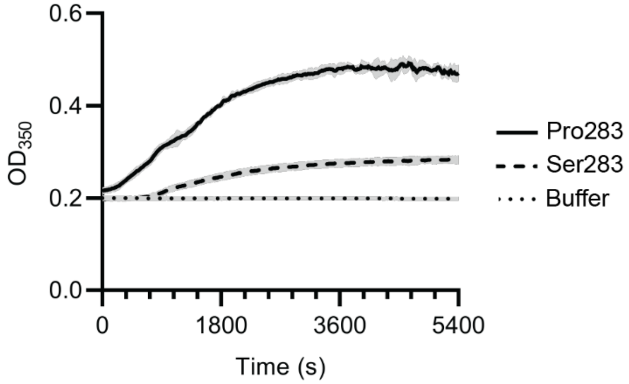


**Fig. S9. Rate of aggregation of NP.**

Denatured NP was refolded at 37 °C, and $OD_{350}$ was plotted against time after mixing. The "Buffer" curve indicates the turbidity change of the refolding buffer in the absence of NP.
